# Prebiotic membrane structures mimic the morphology of purported early traces of life on Earth

**DOI:** 10.1101/2023.11.03.565097

**Authors:** Seán F. Jordan, Mark A. van Zuilen, Joti Rouillard, Zita Martins, Nick Lane

**Author notes:** **Corresponding author:** Seán F. Jordan.

## Abstract

Elucidating the most probable compositions of the first cell membranes prior to the origin of life, within a laboratory setting, requires experiments with organic molecules and chemical conditions representative of those present on the early Earth. As such, the membrane forming molecules used in these experiments are described as ‘prebiotically plausible’, i.e., they could have formed through abiotic reactions and be available for membrane formation prior to the emergence of biology. Similarly, the chemical properties of solutions in which these membranes are formed (e.g., pH, temperature, ionic strength) must represent the early Earth environmental conditions under investigation. Here, using a combined confocal and transmission electron microscopy approach, we show that prebiotically plausible organic molecules, in solutions representative of Hadean submarine alkaline hydrothermal vents, form aggregated structures with substantial morphological diversity. The structures hold the potential for use as traces of prebiotic processes in the ancient rock record. In addition, many of the structures are morphologically similar to those which are presented as early microfossils, thus highlighting the limitations of morphological interpretation in these types of studies. Detailed analyses of abiotic organic structures are essential for our understanding of the earliest living organisms on Earth, as well as for our interpretation of any potential biosignatures recovered in the future from extra-terrestrial bodies.

## 1. Introduction

There are many theories that exist as to how life on Earth arose. Whether in hot springs on land^1^ or hydrothermal vents on the ocean floor^2,3^, all these ideas require an energy source within a geological setting that can fuel chemical reactions. Over time these reactions increased in complexity from geochemistry to organic chemistry, eventually leading to biochemistry with the emergence of life. The gradients within these settings could potentially drive chemical reactions producing organic molecules from inorganic reactants (e.g.,^4–7^). Organic compounds then reacted together forming more complex molecules through increasingly advanced pathways, eventually becoming something like a metabolic pathway: a protometabolism^8^. Without boundaries, this protometabolism may not have been long lived. Products could have quickly become diluted, and reactions could have dissipated giving way to alternate reactions. Like the cell membranes found in all living organisms, some form of compartmentalisation would probably have been required. The cell membranes of all extant living organisms are composed of glycerol phosphate phospholipids^9^. However, these phospholipid membranes are possibly too complex to have formed at the earliest stages of the emergence of life on Earth. Instead, early compartments could have been supplied in the form of vesicles, membrane structures composed of single chain amphiphiles (SCAs) such as fatty acids^10^, giving rise to the first protocells.

It is likely that fatty acids, alcohols, and many other SCAs would have existed in almost any origin of life scenario, and the precursors of membrane-forming molecules may have been synthesised abiotically on the early Earth^11^. The list of prebiotically plausible organics is ever-growing due to results from both laboratory experiments and analysis of real samples^12^. Prebiotic synthesis experiments have achieved the formation of carboxylic acids, amino acids, sugars, and nucleotides^4,5,13–23^. For example, Fischer-Tropsch-Type (FTT) syntheses under hydrothermal conditions produce numerous fatty acids, alcohols, and alkanes containing 6 to 34 carbon atoms^13^, all of which are suitable membrane-forming components. Carboxylic acids and amino acids of abiotic origin have been detected in rock samples from Earth, while numerous organic molecules including both aliphatic and polyaromatic hydrocarbons (PAHs), hydroxy acids, nucleobases, and amino acids have been detected in meteorites^24^. Carbonaceous meteorites contain fatty acids, including aliphatic straight-chain and branched-chain monocarboxylic, with up to 12 carbon atoms, and dicarboxylic acids^25,26^. The straight-chain monocarboxylic acids are dominant, followed by the branched-chain monocarboxylic acids, and finally the dicarboxylic acids. Meteoritic monocarboxylic acids are enriched in deuterium and ^13^C, which is consistent with an extra-terrestrial origin. It has been suggested that gas phase reactions in the interstellar medium, involving radicals and ions, synthesised meteoritic monocarboxylic acids before being accreted in the parent body of meteorites for further processing^27,28^. Carboxylic and dicarboxylic acids can also be formed in the meteorite parent body by two ways: 1) via hydrolysis of carboxamides^29^; or 2) by the oxidation of hydrocarbons from the macromolecular insoluble organic matter (IOM), or the oxidation of free hydrocarbons by oxidised fluids or minerals^30^. Vesicles have in fact been formed directly from organics contained within meteorite samples^31,32^. The meteorite flux to the Earth was much higher during the Hadean than it is today and may have spiked between 4.1 and 3.8 Ga^33,34^. The vast amount of material delivered to the early Earth during this time represents a significant source of organic molecules. Coupled to organics formed *in situ* on the early Earth, it is probable that a wide range of compounds would have been available for membrane formation.

Multiple different early Earth environments have been proposed as possible settings for the emergence of the first living organisms. Each of these scenarios brings with it a unique set of conditions that would potentially affect the production and survival of organic molecules, the formation of membranes and the diversity of possible prebiotic chemical reactions. Deep sea locations with hydrothermal activity could provide a supply of organics through FTT syntheses, with high temperature acidic fluids in ‘black smokers’ and lower temperature alkaline fluids in ‘white smokers’ providing very different pathways for eventual chemical reactions. Some of these sites may have been too deep for meteoritic organic delivery, while being protected from their impacts. Conversely, terrestrial hot springs are exposed to impacts, while simultaneously being receptive to extra-terrestrial organic delivery. Depending on their geology, these surface sites would have unique pH, temperature, and ionic species in their fluids, each combination of which could be amenable to different modes of prebiotic chemistry. These are just two examples of myriad possibilities. It is clear that the permutations of environmental conditions for potential origin of life locations on the early Earth are vast.

Despite this, the majority of work on membrane formation to date has focused on the analysis of vesicles formed from single molecules or simple mixtures containing at most two to three SCAs. The resulting vesicles struggle to survive under environmental stresses such as pH, temperature, and ionic strength fluctuations^35–38^. Sensitivity to salinity, in particular, has cast doubt on any oceanic environment as a possible origin of life location. Recent work, however, has found that SCAs with novel headgroup moieties can withstand some more challenging conditions including high ionic strength and extremes of pH, particularly at acidic levels^39^. It has also been shown that combining simple SCAs in relatively complex mixtures, arguably more relevant to a prebiotic scenario, produces vesicles with significant resilience to multiple environmental stresses, including salinity and high pH^40^. Considering the wide range of available SCAs and the vast amount of research that remains to be done on vesicle formation capabilities, it is clearly unreasonable to exclude any potential origin of life environment based on current knowledge. In fact, it now appears likely that vesicles would have formed in almost any prebiotic scenario.

Vesicles display a diverse range of morphologies, both on an individual level and as aggregates or clusters of multiple vesicles^40,41^. These morphologies seem to be affected by environmental conditions. However, this has not been systematically investigated yet. In fact, the primary objective of origin of life membrane formation studies – testing the ability of membrane formation – has meant that these structures have been overlooked, often regarded as failed experiments. Many of these assemblages are reminiscent in their morphology of living organisms and ancient microfossils. Similar forms created entirely from self-assembled nanocrystalline materials have been described previously^42–45^. It has been suggested that these inorganic ‘biomorphs’ may be observed within the rock record and could be misinterpreted as microfossils of ancient living organisms. The same could be true for organic assemblages of vesicles, and perhaps for biomorphs formed through combinations of both organic and inorganic compounds.

Here we show that combinations of prebiotically plausible organic molecules, in solutions chemically representative of a potential early Earth environment, form a variety of complex aggregate structures. Confocal and electron microscopy reveal the morphological diversity of these structures and the similarities they hold with purported microfossils of the earliest living microorganisms on Earth. We highlight the potential for these abiotic structures as diagnostic signatures in their own right, while also highlighting their tendency to mimic microfossils in the rock record. Population morphometry results from organic biomorphs and microbial populations show strong similarities in 2D structure. However, differences in population size distribution suggest that this characteristic may be useful in distinguishing between abiotic and biological microstructures of this type. Caution is required when investigating signatures of this nature and significantly more work is required to understand these abiotic structures sufficiently, to ensure that they do not hinder our interpretation of biosignatures both on Earth, and potentially elsewhere in our Solar System in the future.

## 2. Methodology

### 2.1 Materials

All reagents used in this study were of analytical grade (≥ 97%) and were procured from either Sigma Aldrich (Merck, UK) or Acros Organics (UK).

### 2.2 Preparation of vesicle solutions

The vesicle solutions were prepared employing a modified version of the technique described by Monnard and Deamer (2003)^46^. Glass vials were employed for all solution preparations in a dry heating block maintained at a temperature of 70 °C. This temperature corresponds to the conditions expected at Hadean alkaline hydrothermal vents (50 – 100 °C)^47^. The molecules used were fatty acids and 1-alkanols (C_10_ to C_15,_ odd and even), and the isoprenoid molecules geranic acid and geraniol. The lipids were heated and introduced to the solutions in their liquid state. Acid molecules were first added to deionized (DI) H2O to provide the desired final concentration and subjected to vortexing. Subsequently, 1 M NaOH was added until the solution turned transparent, indicating complete deprotonation of the acid. The alcohols were then added, and the solution was vortexed. The pH was adjusted by employing 1 M HCl or 1 M NaOH to achieve the desired final value. The solution was then brought to the desired final volume by adding DI H2O. Immediate analysis of solutions was conducted after their preparation.

For the preparation of vesicles in salt solutions, the H2O was substituted with the desired salt concentration, and the acid and base solutions were substituted with the salt dissolved in 1 M HCl and 1 M NaOH, respectively, to ensure consistent salt concentrations throughout. Anoxic vesicle solutions (H2O, FeCl2, Na2S, FeS, Fe particles) were prepared following the aforementioned method in an anaerobic hood with a 5% H_2_ in N_2_ atmosphere. O_2_ was eliminated by reacting with H_2_ to form H_2_O vapour on a Pd catalyst. O_2_ levels were monitored and maintained at 0 ppm throughout these procedures. FeS solutions were prepared using a 1:1 ratio of FeCl_2_ and Na_2_S. Fe particles were added to the relevant solutions after vesicle formation in H_2_O. All anoxic solutions were sealed with lids and wrapped in parafilm. The seal was broken immediately before imaging.

### 2.3 Confocal microscopy

Confocal microscopy was conducted using a Zeiss LSM-T-PMT 880 instrument coupled to an Airyscan detector. Membranes were visualised using the hydrophobic dye Rhodamine 6G. 0.5 µL of a 100 µM dye solution was added to a heated (70 °C) microscope slide. The vesicle solution was vortexed, and a 5 µL aliquot was added to the slide, mixed with the dye. The aliquot was covered with a #1.5, 16 mm diameter coverslip and positioned on the microscope stage. The dye was excited using an Ar laser operating at 514 nm and observed through a 63x oil objective with a 488 nm filter. Images were captured using Zeiss Zen microscopy software, and final processing was conducted using the FIJI (Fiji is just ImageJ) software package.

### 2.4 Negative Staining – Transmission Electron Microscopy (NS-TEM)

NS-TEM was performed using a JEOL 1010 TEM (JEOL, Japan). Samples were applied to a Cu 100 mesh grid and allowed to incubate for 30 seconds. Excess sample was then removed by blotting with filter paper, and a portion of aqueous uranyl acetate (1.5%) was added to the grid. After standing for 30 seconds, the excess liquid was blotted. Grids were immediately analysed under vacuum. Image processing was carried out using the FIJI software package.

### 2.5 Cryo-electron microscopy (Cryo-EM)

Samples were prepared using a Vitrobot Mark IV (Thermo Fisher). An aliquot was applied to a glow-discharged Lacey Carbon (400 mesh Cu) grid (Agar Scientific) for 30 s, blotted for 8.5 or 11 s at 4.5 °C and 95% humidity, and then rapidly plunged into liquid ethane. Imaging was performed on a T10 microscope (FEI) at 100 kV. Images were collected at a magnification range of 7000-34000x.

### 2.5 Population morphometry

Population morphometry analysis^43,48,49^ was performed on several images of the experiments, following the protocols described in Rouillard et al. (2020)^49^. All image treatment was carried out using FIJI version 1.53t. Some improvements in image segmentation were implemented, relative to Rouillard et al. (2020)^49^, particularly making use of machine learning trainable segmentation and threshold binarization. The individual steps of morphometric analysis are described below:

#### 2.5.1 Image segmentation

Using FIJI, each image was cropped to exclude scale bars and other annotated features. For some images with uneven background shading, a subtraction of background noise was performed using the Subtract Background process (using a rolling ball radius of approximately the size of the particles). Further optimalisation before segmentation depended on the individual image. This included image inversion (Fig. S1b), an additional Gaussian blur to make cells stand out relative to artifacts (Fig. S2b), or edge-detection (using the Canny Edge plugin) followed by hole-filling (Fig. S3c).

Subsequently the Trainable Weka Segmentation plugin^50^ was loaded. The classifier training process was initiated using two classes, one for the structures of interest and another for background (Fig. 1; Fig. S1 c, d; Fig. S2 c, d). The classifier was trained to fill holes and to separate touching cells. In some images it was more effective to classify artifacts that obscured the cells (Fig. 3; Fig. S3c-f). In that case an image was created that could be subtracted from the pre-Weka image, resulting in a cleaned-up version showing only the particles of interest. For all Weka-segmented images a binarized segmented image was then created (Fig. S1e, S2e, S3g). Individual particles could then be counted using the Analyze Particles option, selecting area and shape descriptors. The major steps of this protocol are shown in Figure 1.

**Figure 1.**
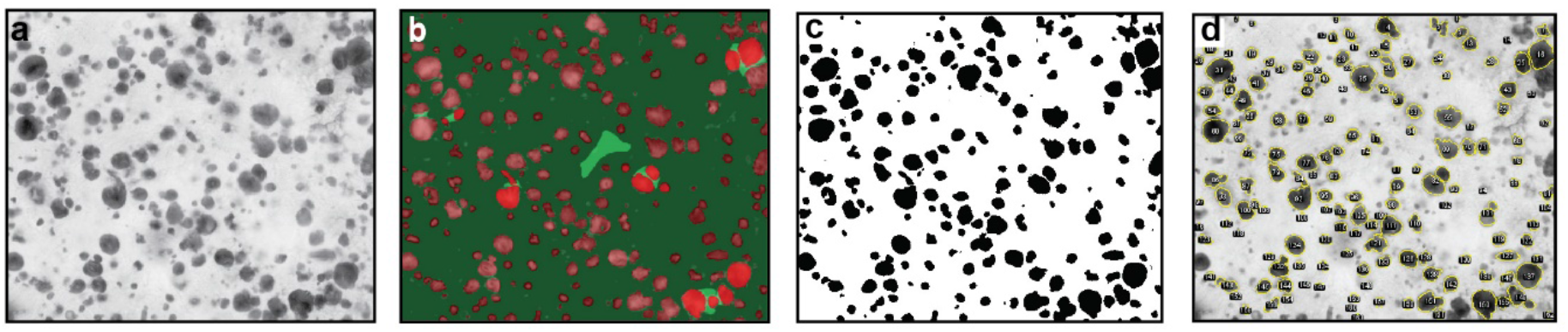
Major steps in image segmentation and particle counting protocol. Experimental vesicles (dried). a) Original, cropped image, b) Weka trainable segmentation, separating individual cells from background, c) thresholding and binarization, d) Particle counting, size and shape description. For more details on segmentation, see Figure S1.

#### 2.5.2 Particle characterization

Following Rouillard et al. (2020)^48^, for each segmented image the entire population of particles was counted (using a 5-pixel size threshold to avoid small dots), and for each particle the area (A) (in square μm), radius (R) (assuming a perfect circle, in μm) circularity (C) and solidity (S) were determined. Circularity is defined as C = 4πA/P2, where P (in pixels) is the circumference of the particle. Solidity is defined as S = A/α where α is the area (in square pixels) within the convex hull of the particle. This convex hull consists of the surface bound by straight lines that join the outermost points of the particle.

## Results

A selection of prebiotically plausible and biochemically important single chain amphiphilic organic molecules were used to investigate the formation of abiotic structures (Table 1). Different combinations of organics, pH and additional ionic species were used for each solution prepared. Molecules were first dissolved in alkaline (pH ca. 11) aqueous NaOH solutions. This pH is sufficiently above the pKa of the individual molecules to provide solutions so that the organics exist as monomers or micelles. The solutions were titrated with 1 M HCl to encourage bilayer formation leading to the development of individual vesicles and more complex structural assemblages. Confocal micrographs of ‘typical’ solutions present with multiple circular and elliptical vesicles (Fig. 2a-c). Some vesicles can become elongated and others encapsulated within larger vesicles (Fig. 2b and c). Vesicles are generally floating in solution unobstructed during analysis and display some flow caused by capillary action on the glass slide and erratic movement likely due to Brownian motion (SI video). TEM is performed under vacuum and as such the resulting micrographs contain collapsed or ‘doughnut’ shaped vesicles (Fig. 2d-f). These would have been spherical in shape prior to being exposed to the vacuum pressure.

**Table 1.**
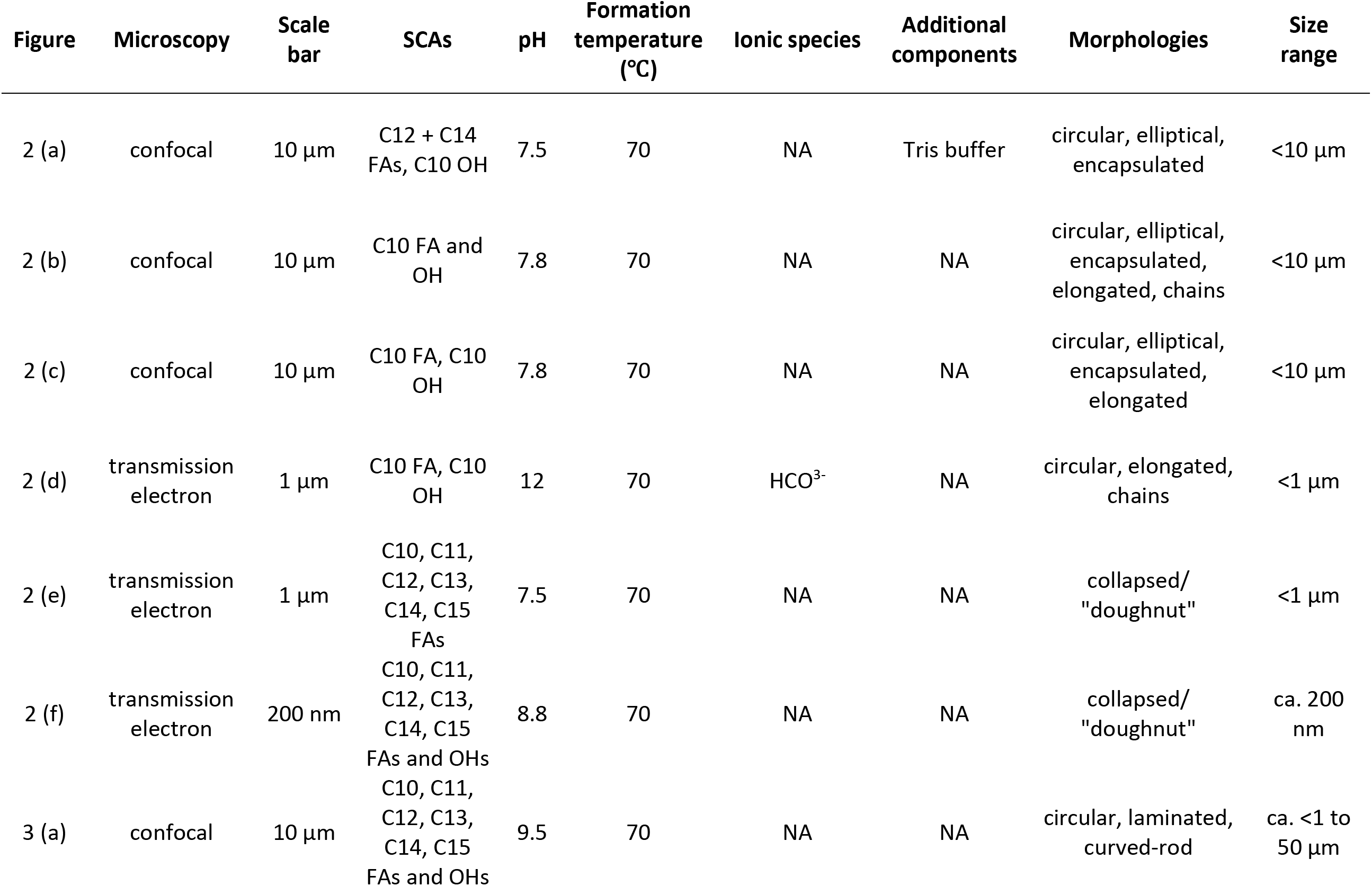
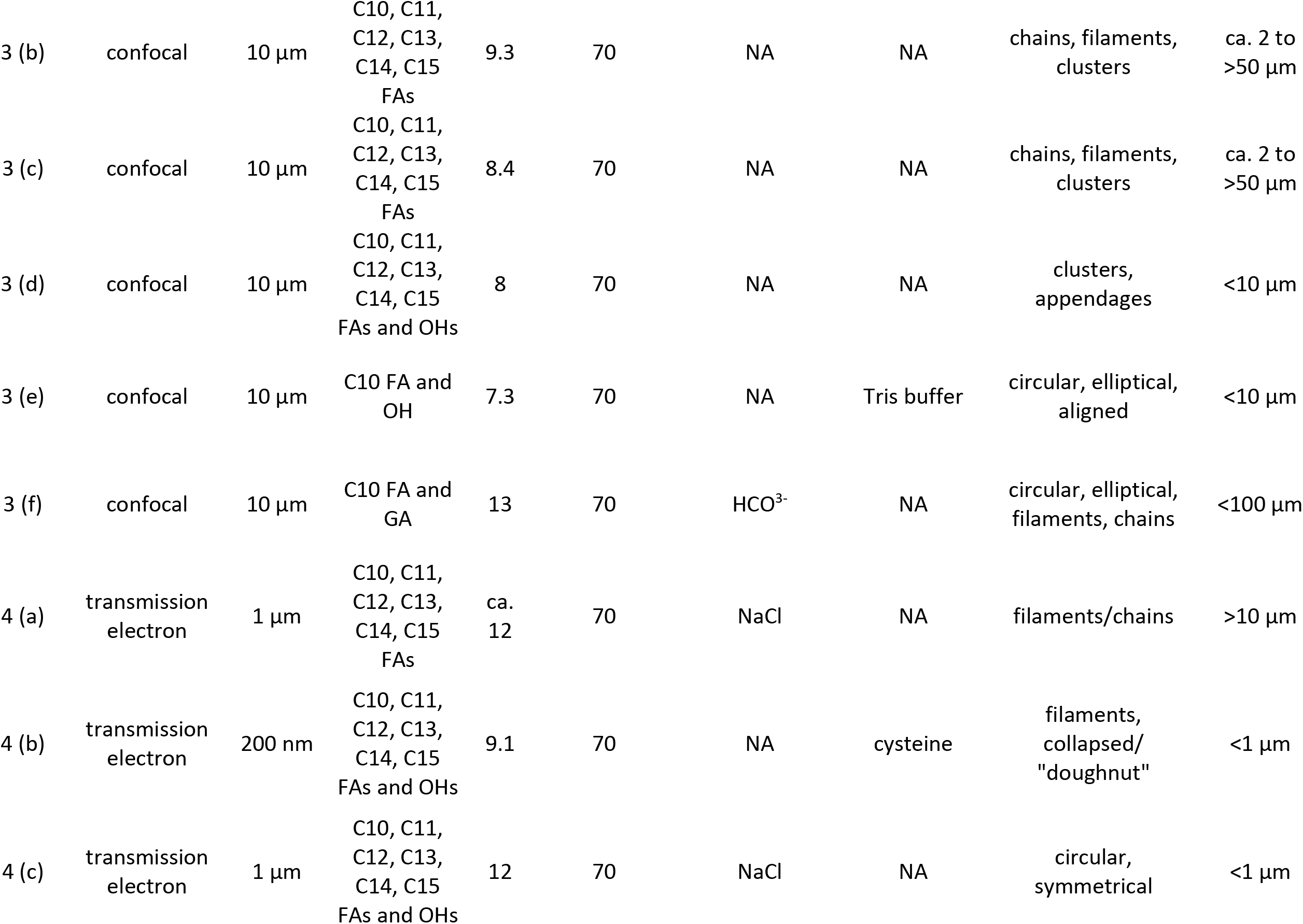
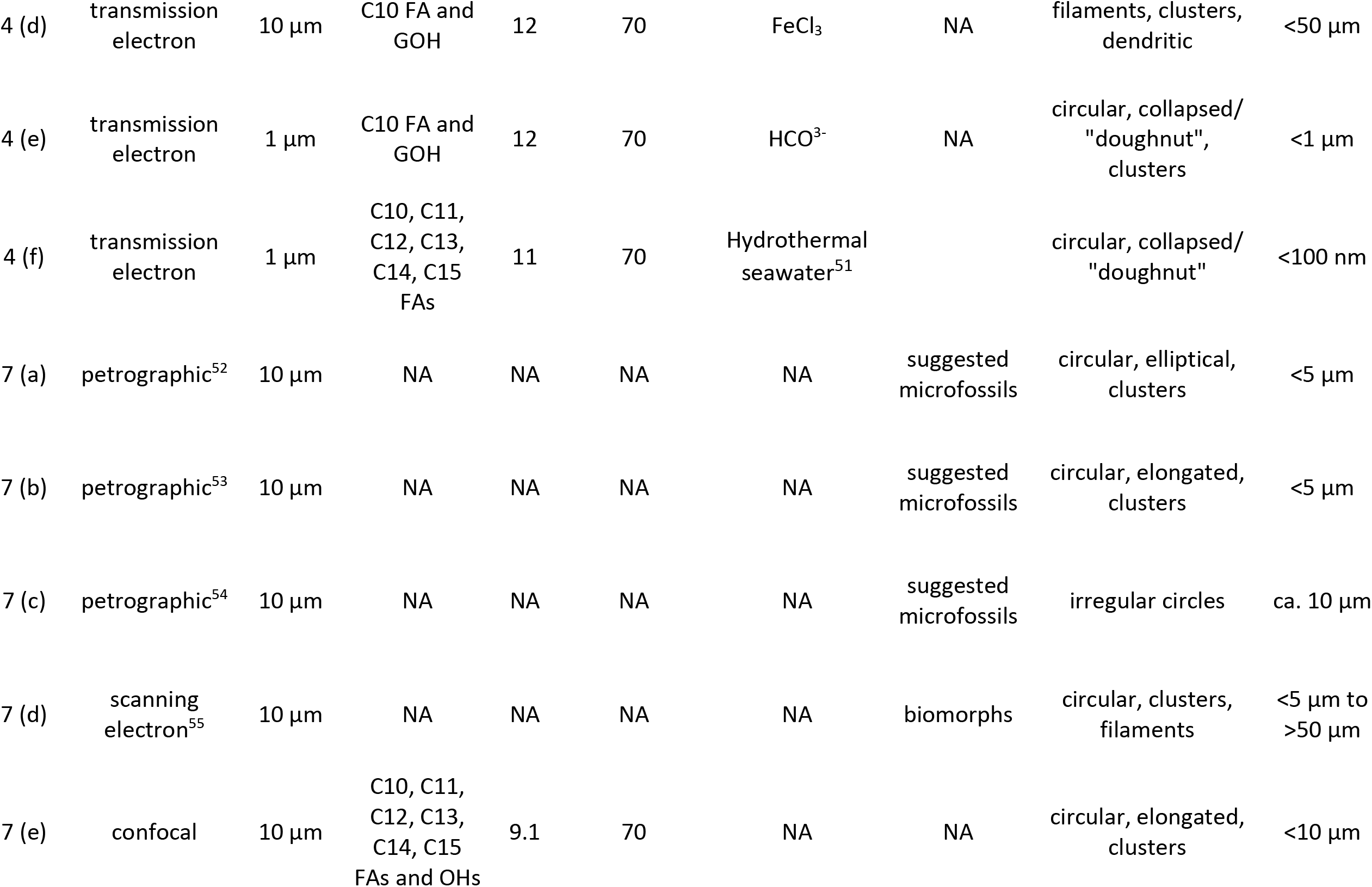
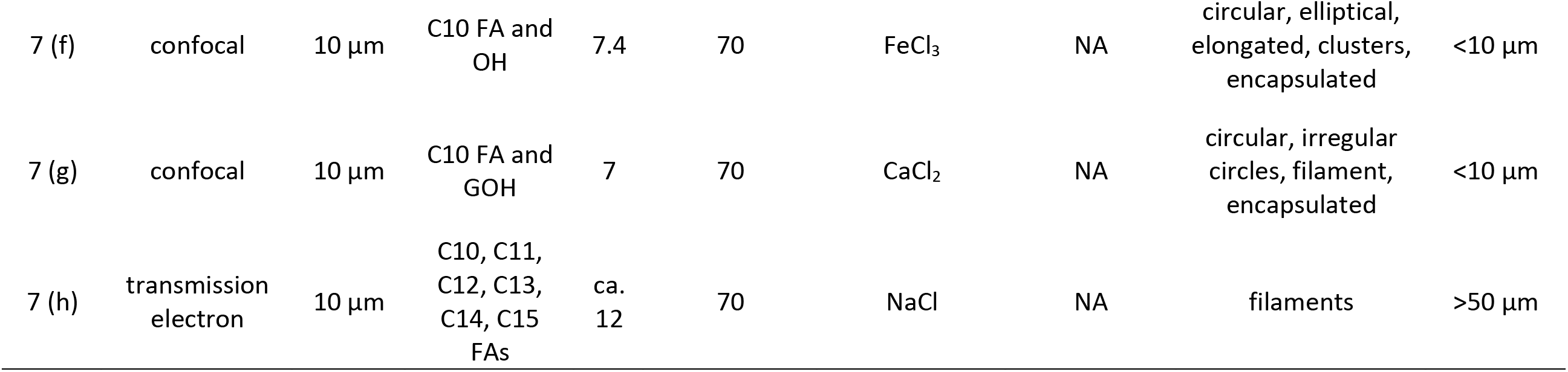
Full details for each figure panel. SCA – single-chain amphiphile; FA – fatty acid; OH – alcohol; GOH – geraniol.

**Figure 2.**
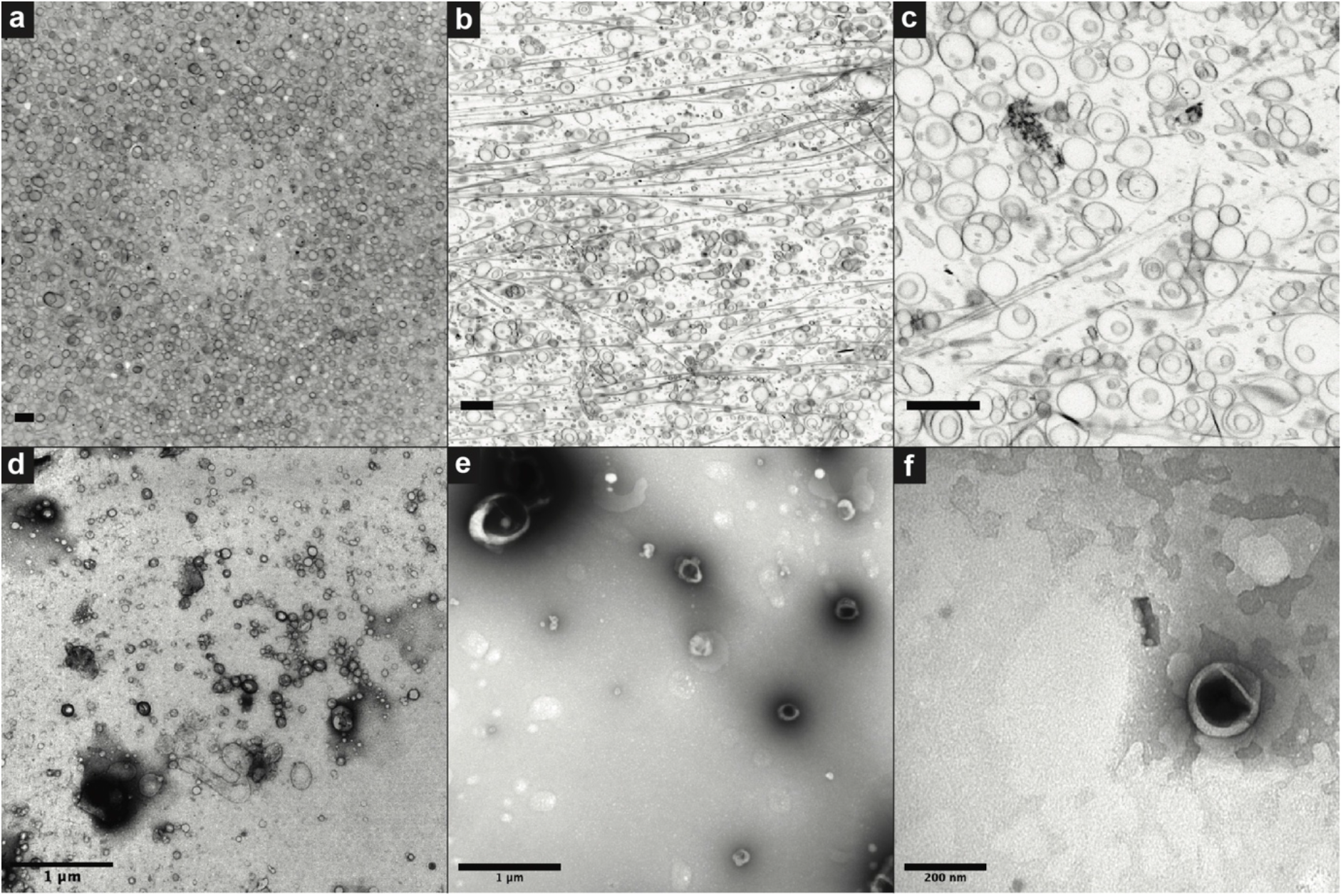
Micrographs of images from ‘successful’ vesicle formation experiments. (a-c) Vesicles in solution imaged using confocal microscopy, (d-f) transmission electron micrographs of vesicles mounted on Cu grids. Scale bars are 10 µm (a-c), 1 µm (d and e) and 200 nm (f).

As fluid flows on the glass slide during confocal analysis, some vesicles remain in place adhering to the glass and subsequently drying out. This can leave traces in a variety of shapes including laminated, curved-rods (Fig. 3a), large clusters with appendages (Fig. 3d), and filled circles or ellipses connected to filaments (Fig. 3f). Filaments are a common occurrence in solution (Fig. 3b and c) although their formation mechanism is unclear. Individual vesicles can also settle in formation while still in solution (e.g. linear formation in fig. 3e). This is likely due to a physical deformation in the glass slide which is linear in shape or the presence of a linear contaminant object to which the vesicles are attracted, potentially by surface charge interactions. Further work is necessary to elucidate the exact mechanisms behind these formations.

**Figure 3.**
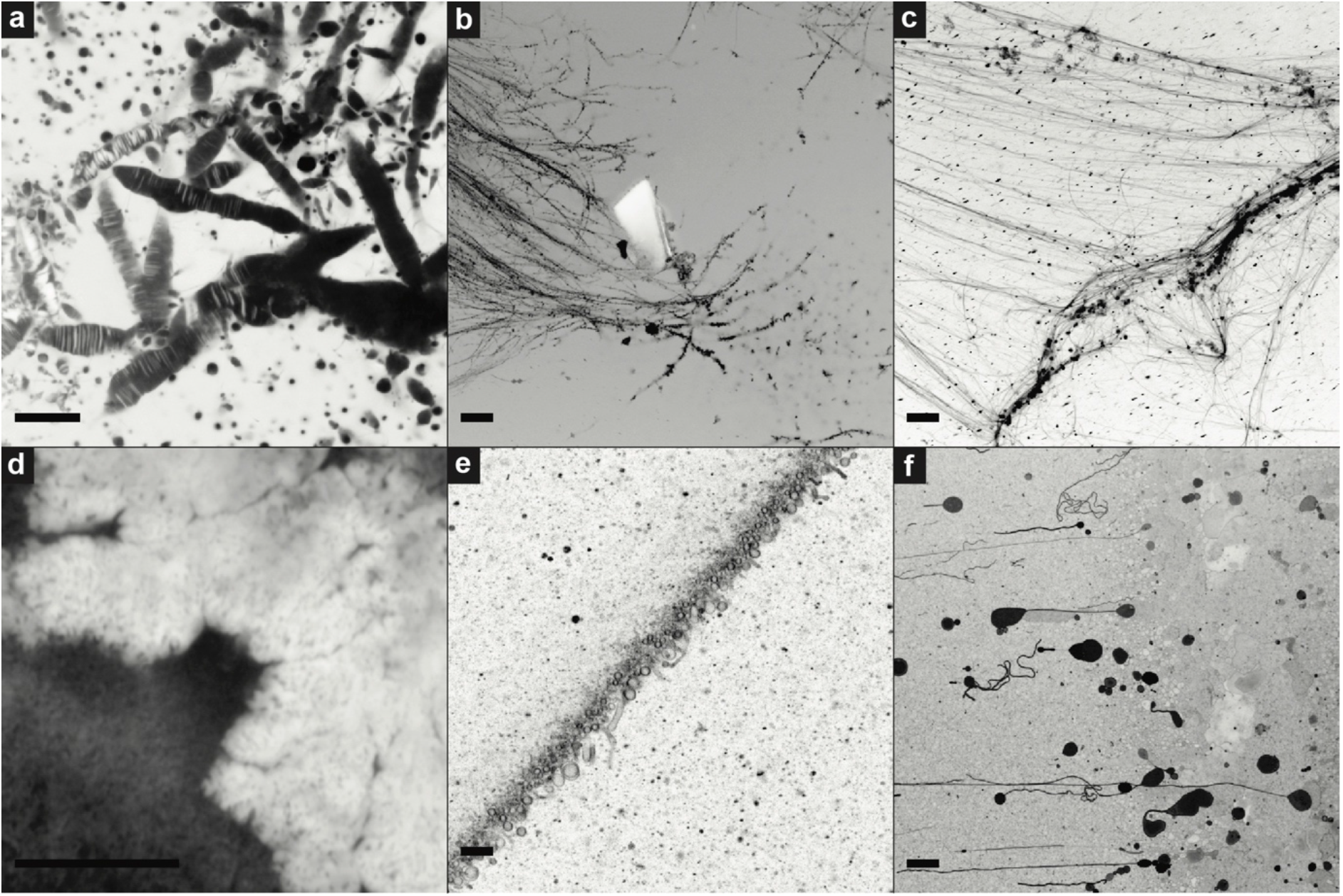
Confocal micrographs of aggregates formed from solutions of organic molecules used in vesicle formation experiments. All scale bars are 10 µm.

The diversity of assemblage morphologies increases when analysed by TEM (Fig. 4). The vacuum clearly plays an important role here as in the drying during confocal analysis which also produced unique structures. Collapsed or ‘doughnut’ structures remain common but are often accompanied by filaments, chains, dendritic, and symmetrical patterns. The cause of these patterns is unclear although the presence of inorganic components in the solutions tends to increase the likelihood of complex morphologies. Some of the filamentous structures are comprised of individual vesicles and have been observed previously during confocal microscopy and subsequent TEM of the same solutions^40^. It was suggested that these filaments are the result of the organic vesicles being essentially salted-out of solution by the presence of inorganic salts such as NaCl.

**Figure 4.**
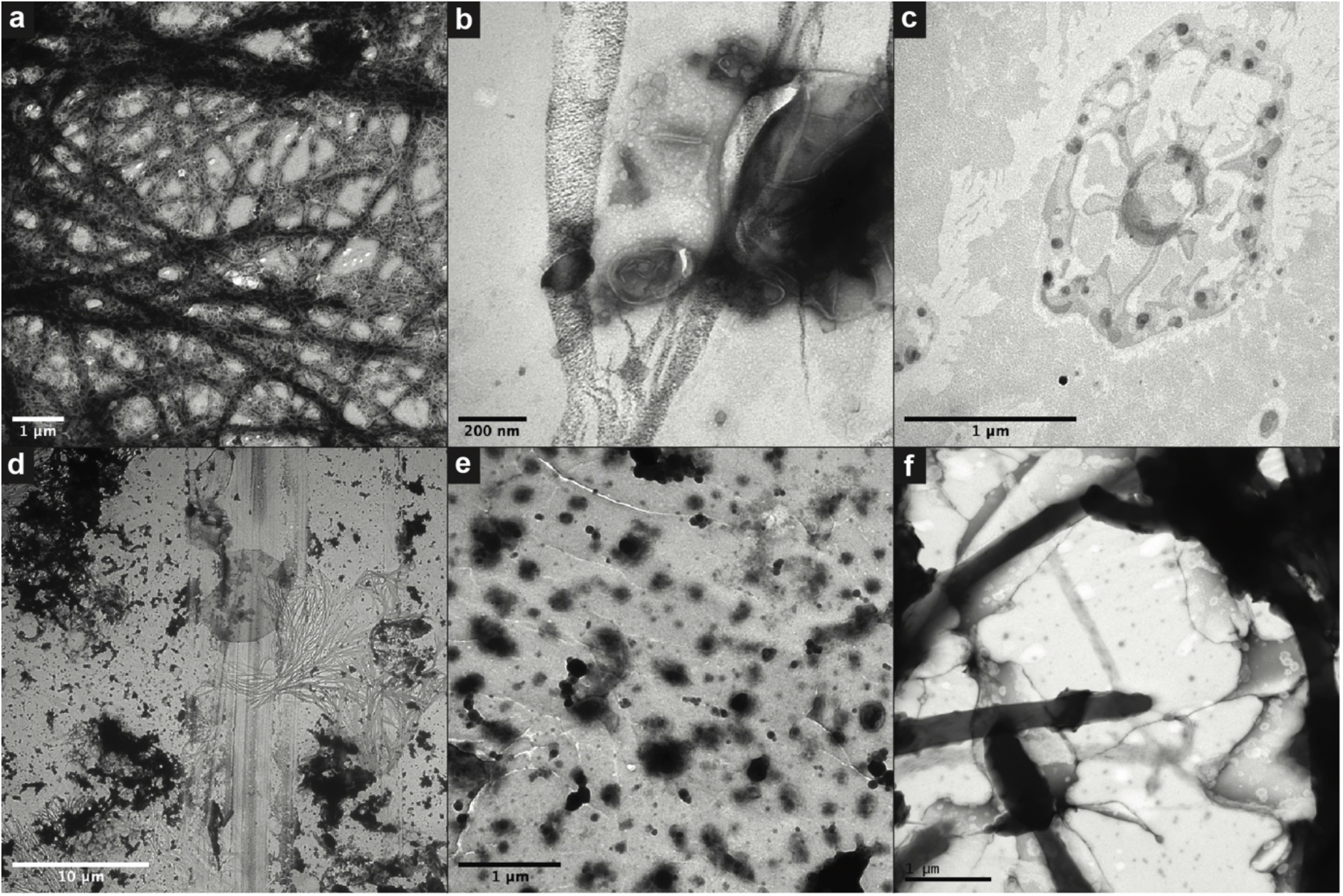
Transmission electron micrographs of aggregates formed from solutions of organic molecules used in vesicle formation experiments. Scale bars are 1 µm (a, c, e and f), 200 nm (b) and 10 µm (d).

The population morphometry results for experimental vesicles were compared with a population of coccoidal cyanobacteria (Fig. 3a in Ref^48^,). For this latter image the exact same (updated) segmentation protocol was used (Fig. 5). The results are shown in Table 2, Fig. 6 and Fig. S4. Both circularity (C) and solidity (S) values for biomorph images (C = 0.83 – 0.84, S = 0.90 – 0.95) are similar to those of the cyanobacteria (C = 0.88, S = 0.96). The primary difference between biomorphs and microorganism in these analyses lies in the size measurement, where the cyanobacteria have a narrow size distribution (Fig. 6c, Fig. S4, radius mean/Std = 5.37) while the biomorphs have a radius mean/Std = 2.33 and radius mean/Std = 1.55 for dried and cryo experiments, respectively (Fig. 6b, c; Fig. S4). Overall, these data show that these organic biomorphs have a less uniform size distribution than typical single-strain populations of microbial cells, but that their shape may be indistinguishable. Given that a single parameter is not a reliable indicator, further experimentation is needed to expand our understanding of population morphometry in this area. Using a broader range of biomorphs, microbial populations, and microfossils will allow us to determine if there are statistically significant differences that will enable this approach to be used to determine biogenicity in the future.

**Figure 5.**
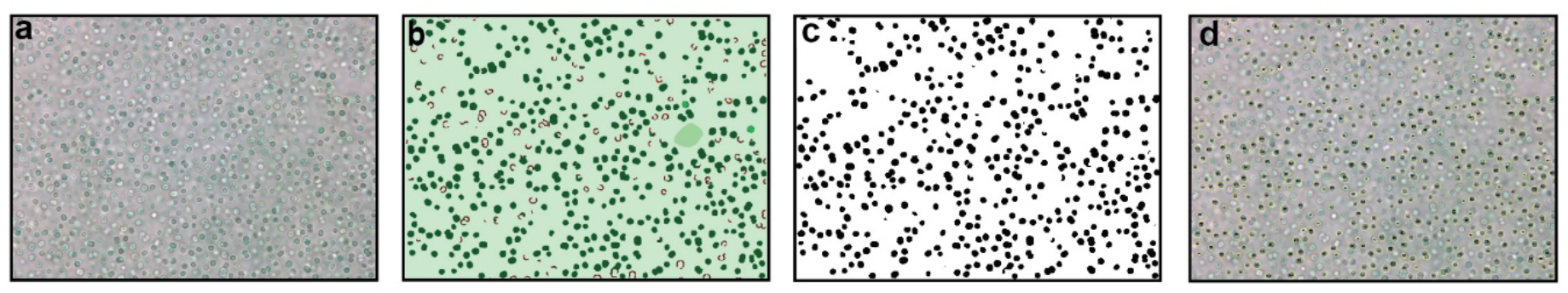
Image treatment for Synechocystis sp. (PCC6803), following the same protocol as for the experimental vesicles, shown in Fig. 1a) Original, cropped image, b) Weka trainable segmentation, separating artifacts from cells and background (see Fig. S3 for details), c) Tresholding and binarization, d) Particle counting, size and shape description. For more details on segmentation see Figure S3.

**Table 2.**
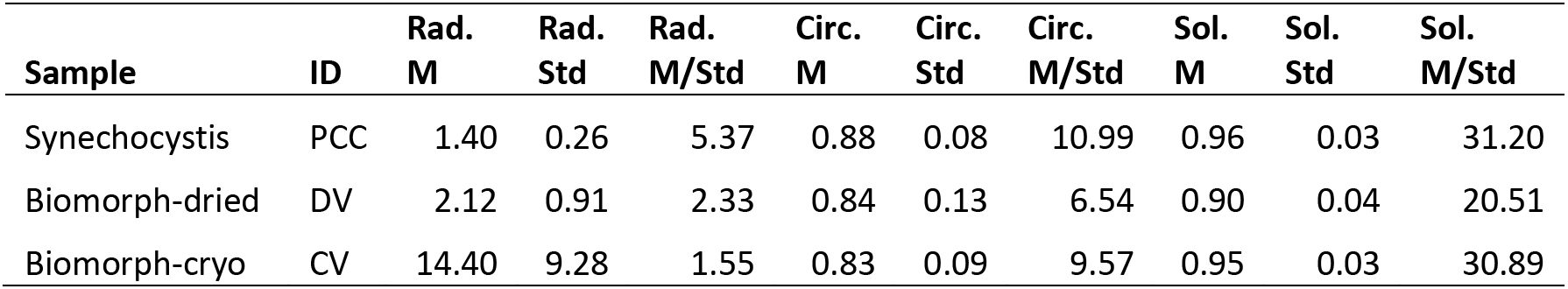
Particle characterisation parameters for Synechocystis sp. and biomorphs.

| Sample | ID | Rad.<br>M | Rad.<br>Std | Rad.<br>M/Std | Circ.<br>M | Circ.<br>Std | Circ.<br>M/Std | Sol.<br>M | Sol.<br>Std | Sol.<br>M/Std |
| --- | --- | --- | --- | --- | --- | --- | --- | --- | --- | --- |
| Synechocystis | PCC | 1.40 | 0.26 | 5.37 | 0.88 | 0.08 | 10.99 | 0.96 | 0.03 | 31.20 |
| Biomorph-dried | DV | 2.12 | 0.91 | 2.33 | 0.84 | 0.13 | 6.54 | 0.90 | 0.04 | 20.51 |
| Biomorph-cryo | CV | 14.40 | 9.28 | 1.55 | 0.83 | 0.09 | 9.57 | 0.95 | 0.03 | 30.89 |

**Figure 6.**
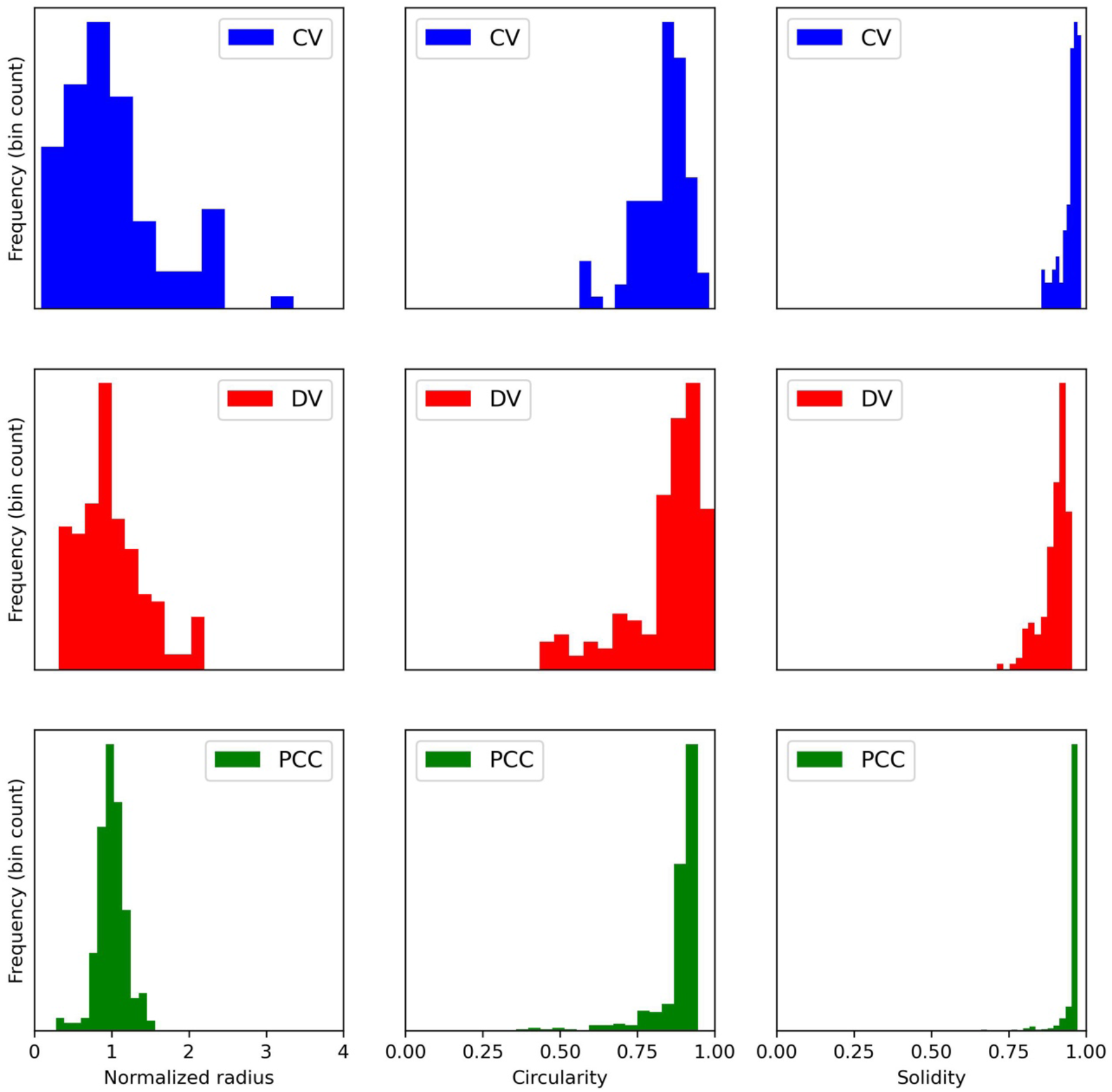
Distributions of normalized size, circularity and solidity of cryogenic vesicles (Fig. S2), dried vesicles (Fig. S1) and microbial cells (Synechocystis sp., PCC6803, Fig. S3).

## Discussion

Our results show that abiotic organic molecules, likely present on the early Earth, including fatty acids, 1-alkanols, and isoprenoids are capable of forming a diverse range of complex morphologies. Confocal microscopy and TEM analyses provide insights into the possible forms observed using different analytical techniques. The microstructures are formed across a range of environmental conditions with varying pH, temperature, and ionic strength – each of which appear to affect structural morphology. The vast possible combinations of abiotic organic molecules, inorganic materials, and chemical conditions on the early Earth suggest that the formation of these kinds of biomorphs would have been inevitable. Yet our understanding of their formation mechanisms and diagnostic characteristics remains in its infancy, primarily due to a paucity of research in this area.

### Comparison with microfossils

From the relatively small selection of abiotic mixtures presented here, a substantially diverse range of morphologies was observed some of which are reminiscent of suggested microfossils from Earth’s early rock record (Fig. 7). Vesicle experiments were not directed in any way towards the replication of these or any microfossil morphologies. Yet, many microfossil morphologies reported in the Archean and Paleoproterozoic rock record appear to have comparable shapes and sizes. Clusters of carbonaceous spheroids and filaments, interpreted as ancient cell colonies, have been reported in e.g. 3.4-3.2 Ga cherts of the Barberton Greenstone Belt (BGB), South Africa^52,56–58^, 3.5-3.0 Ga cherts of the Pilbara Granitoid-Greenstone Belt, Western Australia^53,54,59–64^, and the 1.9 Ga Gunflint Formation, Canada^53,65,66^. Some examples of morphological comparison between microfossils and experimentally-produced vesicles are shown in Figure 7.

**Figure 7.**
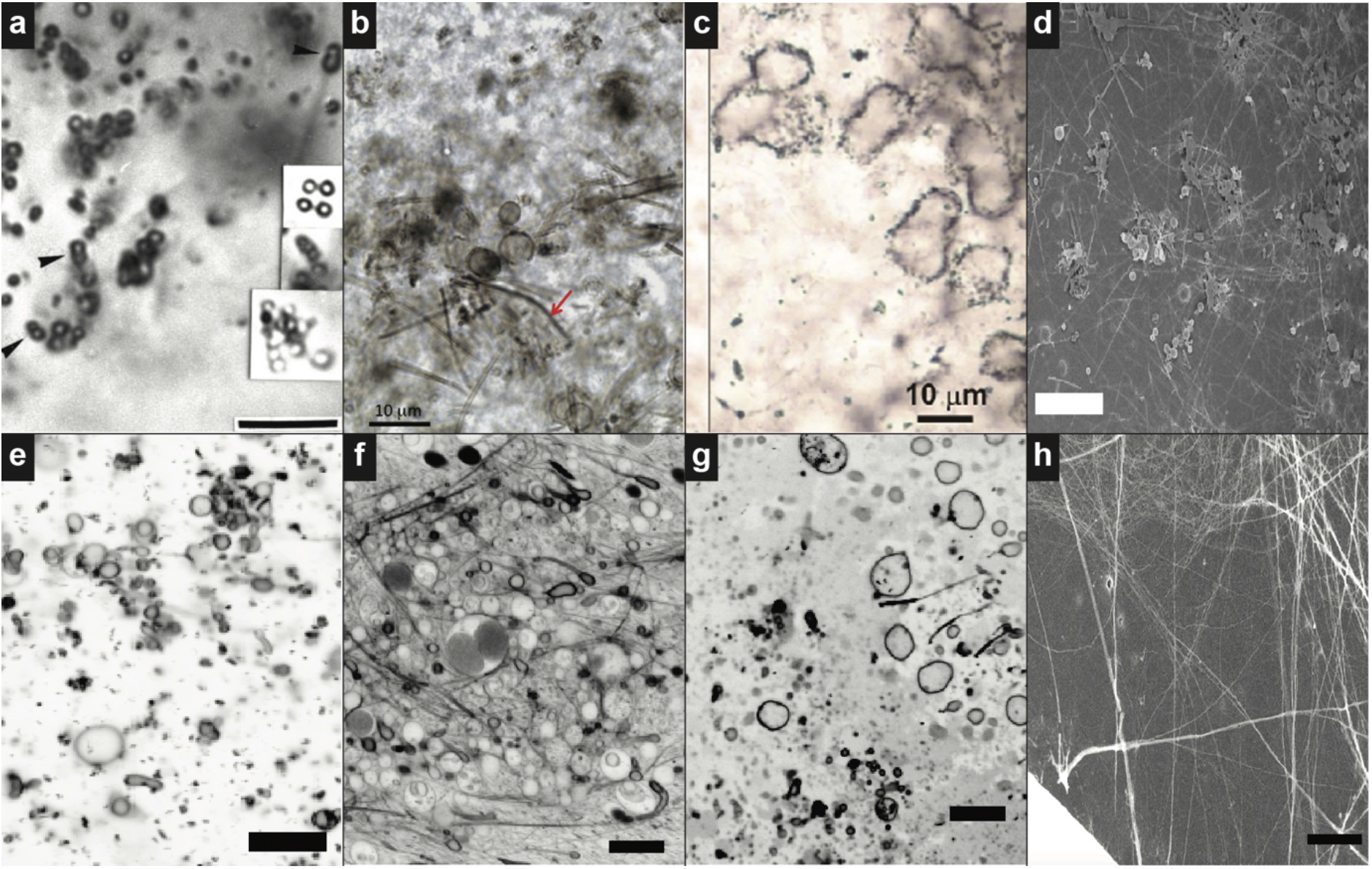
Comparison between purported microfossils and vesicle aggregates. Panels a-c are published micrographs of controversial microfossils (a - Westall et al. (2001), b - Wacey et al. (2012), c - Wacey et al. (2011)), d is a published micrograph of a biomorph from Cosmidis and Templeton (2016) formed from sulfur and yeast extract, and panels e-h are micrographs of vesicles that were produced during this study. All scale bars are 10 µm.

For instance, spheroidal or spherule-like microfossils in the ca. 3.3-3.4 Ga Kromberg Formation within the BGB (Fig. 7a)^52^, are similar in shape to many abiotic vesicle micrographs including the example presented in Figure 5e which were formed from a mix of twelve fatty acids and 1-alkanols (C10 to C15) at alkaline pH (pH 9.1). Westall et al. (2001)^52^ present several examples of spherules ‘joined in pairs’, a feature which is also observed in vesicle solutions. They argue that their observed structures fulfil several biological criteria including size, morphology, cell division, colonial distribution, and cell wall texture. While we agree with this assessment, the vesicle experiments here also arguably meet these criteria. In another example, clusters of spheroidal microfossils, identified in the ca. 3.4 Ga Strelley Pool Formation, Western Australia, and interpreted as remnants of sulphur-metabolising cells^34^ (Fig. 7c), are similar in shape to the ca. 10 µm diameter circular vesicles formed in the laboratory from decanoic and geranic acid in the presence of CaCl_2_ (Fig. 7g).

These morphological similarities extend to filamentous shapes and even to mixtures of spheroids and filaments. For example, Wacey et al. (2012)^53^ reported a selection of spheroidal and filamentous microfossils (suggested Huroniospora and Siphonophycus/Gunflintia respectively) from within a stromatolitic chert of the ca. 1.9 Ga Gunflint Formation, Canada (Fig. 7b). We also observed these co-occurring morphologies in solutions of decanoic acid and 1-decanol containing FeCl_3_ at pH 7.4 (Fig. 7g), although combined circular and filamentous vesicle morphologies are not restricted to Fe-containing solutions. Considering the younger age of these microfossils a biogenic origin is likely but without more advanced data on the formation and preservation of abiotic microstructures it remains uncertain.

It should be noted that, although the vast majority of Archean and Paleoproterozoic microfossil morphologies are spheroidal or filamentous, other morphologies exist such as spindle-like and lenticular shapes^61,67,68^ and hollow tubular organic structures^69^, that do not appear to be mimicked in our current experiments. It may be possible that these morphologies result specifically from the larger molecules which make up prokaryotic cell walls (i.e., polysaccharides and proteins). Indeed, much of the biogenicity criteria employed for microfossil interpretation relates to interpreting cell wall morphology specifically. Further work is needed to elucidate this possibility. That lipid vesicles are chemically analogous to cell membranes rather than cell walls should not however affect their relevance to microfossil interpretation. In chemical terms, following billions of years of diagenesis, organic molecules will largely have been converted to graphite regardless of the complexity of their original form. Some of the biomorph morphologies presented here result from individual vesicles forming individual spherical shapes (e.g., Fig. 2a). Others, such as the filamentous structures, are formed from aggregates of individual vesicles (e.g., Fig. 4a). In addition, it has been shown previously that vesicles can bind to and coat mineral particles, likely due to surface charge interactions, creating an organic cast of the mineral shape^41^. Self-assembly of abiotic SCAs may yet yield these additional morphologies, particularly in conjunction with relevant inorganic species. Moreover, detailed physical and chemical analyses of organic biomorphs will allow us to better understand their relevance to microfossil interpretation.

It is important to realize that up to this point only the morphological aspects of these abiogenic vesicles have been studied and compared to ancient microfossils. In recent years, however, it has become possible to image and characterize individual microfossils using in-situ analytical techniques such as Raman spectroscopy, confocal laser scanning microscopy (CLSM), Secondary ion mass spectrometry (SIMS, NanoSIMS), transmission electron microscopy and energy-dispersive X-ray spectroscopy (TEM-EDXS), and synchrotron-based scanning transmission X-ray spectroscopy (STXM). Combinations of these techniques make it possible to directly link chemical and isotopic characteristics to the morphology of microfossils. A recent overview of these techniques for the study of microfossils is given in Lepot (2020)^70^. It is thus important to note that some of the examples of microfossils given above have been studied using additional chemical and isotopic analyses. For the spheroidal microfossils identified by Wacey et al. (2011)^54^ in the ca. 3.4 Ga Strelley Pool Formation, Western Australia (Fig. 7c), specific SIMS-based δ^13^C and δ^34^S analyses provided evidence for sulphur-based metabolism. For the filamentous and spheroidal microfossils of the 1.9 Ga Gunflint Formation Wacey et al. (2012)^53^ (Fig. 7b) performed more advanced morphological and elemental analyses utilising focused ion beam (FIB) milling in combination with SEM and TEM to provide micrographs of both nanoscale resolution and in three dimensions. In general, a range of chemical-and isotopic biogenicity criteria have thus been created that can be directly linked to the classical morphology-based biogenicity criteria. To date, this analytical approach has not been applied to abiotic vesicle experiments although it is clearly necessary to provide control data for these kinds of microfossil investigations.

### Comparison with other organic biomorphs

Previous work from Cosmidis and Templeton^55^ investigating the potential formation of carbon/sulfur-based biomorphs produced a variety of spheroidal and filamentous structures (Fig. 7d) which resemble the morphologies of filamentous vesicle solutions (Fig. 7h). This highlights the potential for abiotic organic compounds from multiple sources to form microstructures that mimic microfossils. The authors in this case call for ‘new caution’ in interpreting putative microfossils in the rock record^55^. In a recent study Nims et al.^71^ produced biomorphs containing sulphur and organic carbon which were morphologically reminiscent of the filamentous sulphur-oxidizing bacteria *Thiothrix* containing intracellular sulphur globules. Future isotopic analyses performed on these biomorphs could provide some initial control data for sulphur and carbon containing microfossil interpretation. More experimental work of this kind is imperative for our interpretation of microstructures in the rock record. In another recent study Criouet et al. (2021)^72^ showed experimentally that under diagenetic conditions RNA molecules mixed with quartz and water will create spheroidal organic microstructures that mimic microorganisms such as Staphylococcus or Thermococcales. With these types of studies, we are beginning to see only the tip of the iceberg of the potential for organic biomorphs to obscure the rock record.

### Relevance for biogenicity tests

Recently it has been suggested that the development of a strong set of criteria for determining the biogenicity of microfossils is crucial to our understanding of the Earth’s early biosphere^73^. Clearly the design of such criteria necessitates knowledge of the potential for abiotic structures to obscure our interpretations. These microstructures may provide answers to questions that currently overshadow uninterpreted signatures which have been observed in rock samples. It is essential that we understand the formation and preservation of these structures particularly following diagenetic processes representative of those experienced by relevant geological materials. Without the requisite control data, we may continue to struggle through ambiguity. Understanding the composition of these non-living structures would not only improve our interpretation of potential microfossils, but it could also shed light on the environment in which abiotic structures were formed, further adding to our knowledge of Earth’s history. Furthermore, these biomorphs may be particularly beneficial for the analysis of potential biosignatures on other planets where life may have started but perhaps did not evolve in complexity. This is critical for the development of strategies for life-detection and instruments for future space missions^73–75^. With sample return from Mars likely to be realised in the not too distant future, it is essential that we are prepared to provide the best possible interpretation of any proposed biosignatures that we identify during investigations here on Earth.

## Conclusions

The current study presents a direct link between the origin of life and microfossil interpretation. We have shown that lipid vesicles, representative of the first stages of cell membrane development at the emergence of life, are capable of satisfying morphology criteria applied to traces of life from the early Earth, and the development of advanced statistical approaches is required to pinpoint potential differences which may enable us to distinguish between biogenic and abiogenic structures. It has been suggested that the preserved remnants of prebiotic chemistry could potentially satisfy many of the widely accepted biogenicity criteria currently applied to possible microfossils^75^. The work presented here represents a miniscule fragment of all possible prebiotic biomorphs which could be observed in geological samples from the early Earth and potentially elsewhere in the Solar System. These biomorphs could also inform on potential fossilised signatures of life’s emergence which could be observed in either terrestrial or extra-terrestrial samples – protobiosignatures. A significant effort is required to determine the full range of possible prebiotic biomorphs which could affect our interpretation of the earliest rock record including advanced physical, chemical, and statistical analytical approaches.

## Author contributions

Conceptualisation, S.F.J. and M.A.vZ.; Methodology, S.F.J, M.A.vZ., and J.R.; Experiments and analysis, S.F.J., M.A.vZ., and J.R.; Writing—original draft, S.F.J.; Writing—review & editing, S.F.J., M.A.vZ., J.R., Z.M., and N.L.; Project administration, S.F.J.; Funding acquisition, S.F.J, M.A.vZ., Z.M., and N.L.. All authors have read and agreed to the published version of the manuscript.

## Acknowledgements

We thank Mark Turmaine and Andrew M. Hartley for assistance with NS-TEM and Cryo-EM experiments, respectively. S.F.J. acknowledges support from ”la Caixa” Foundation (ID 100010434) and from the European Union’s Horizon 2020 research and innovation programme under the Marie Skłodowska Curie grant agreement No 847648. The fellowship code is “LCF/BQ/PI21/11830015”. Centro de Química Estrutural acknowledges the financial support of Fundação para a Ciência e Tecnologia (FCT) through projects UIDB/00100/2020 and UIDP/00100/2020. Institute of Molecular Sciences acknowledges the financial support of FCT through project LA/P/0056/2020. Z.M. acknowledges funding from Fundação para a Ciência a Tecnologia (FCT) grant number 2022.05284.PTDC.

