## Supplementary figures and images for "Prebiotic membrane structures mimic the morphology of purported early traces of life on Earth"

### S1

**Figure S1 Dried vesicles**

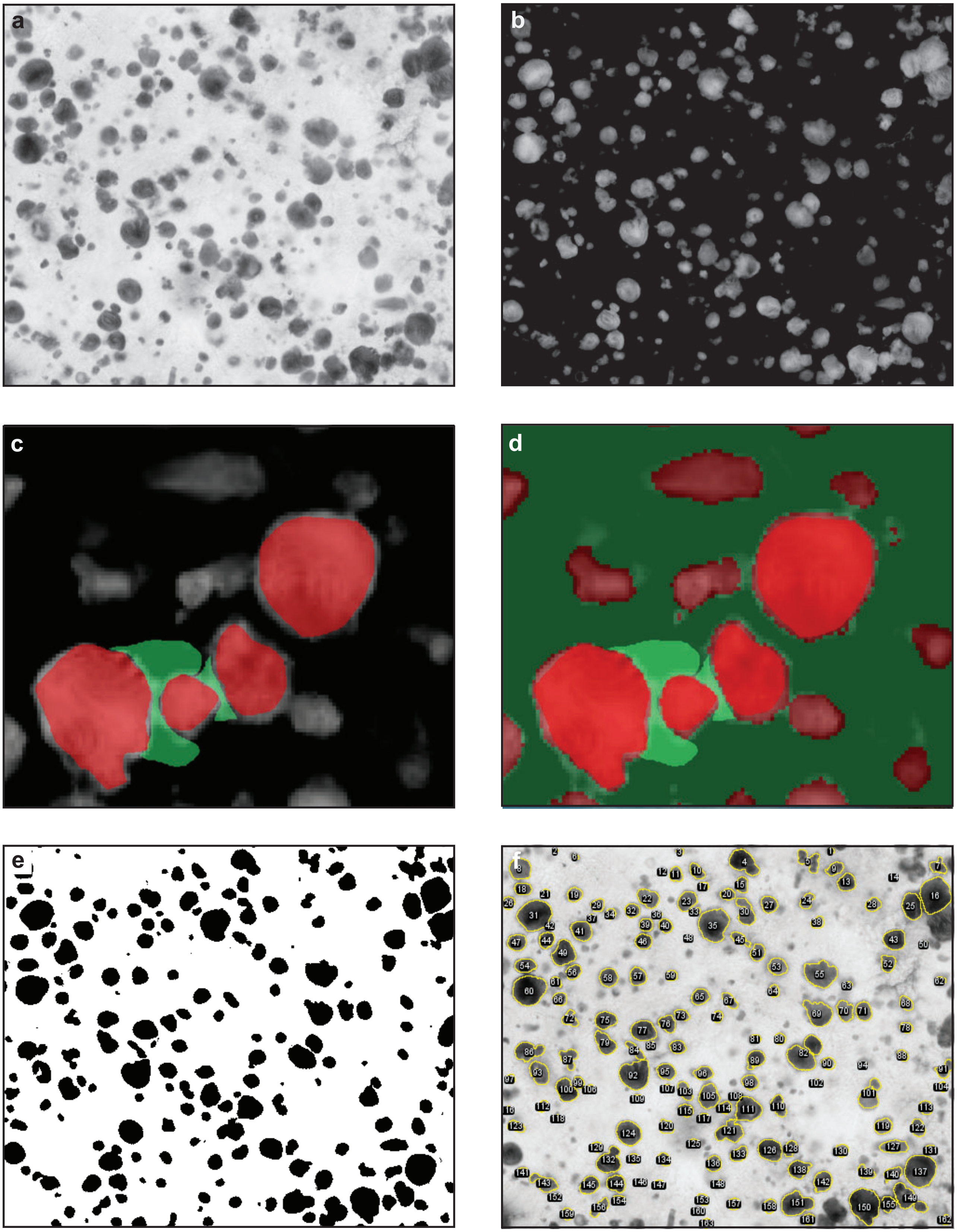

### S2

**Figure S2 Cryo vesicles**

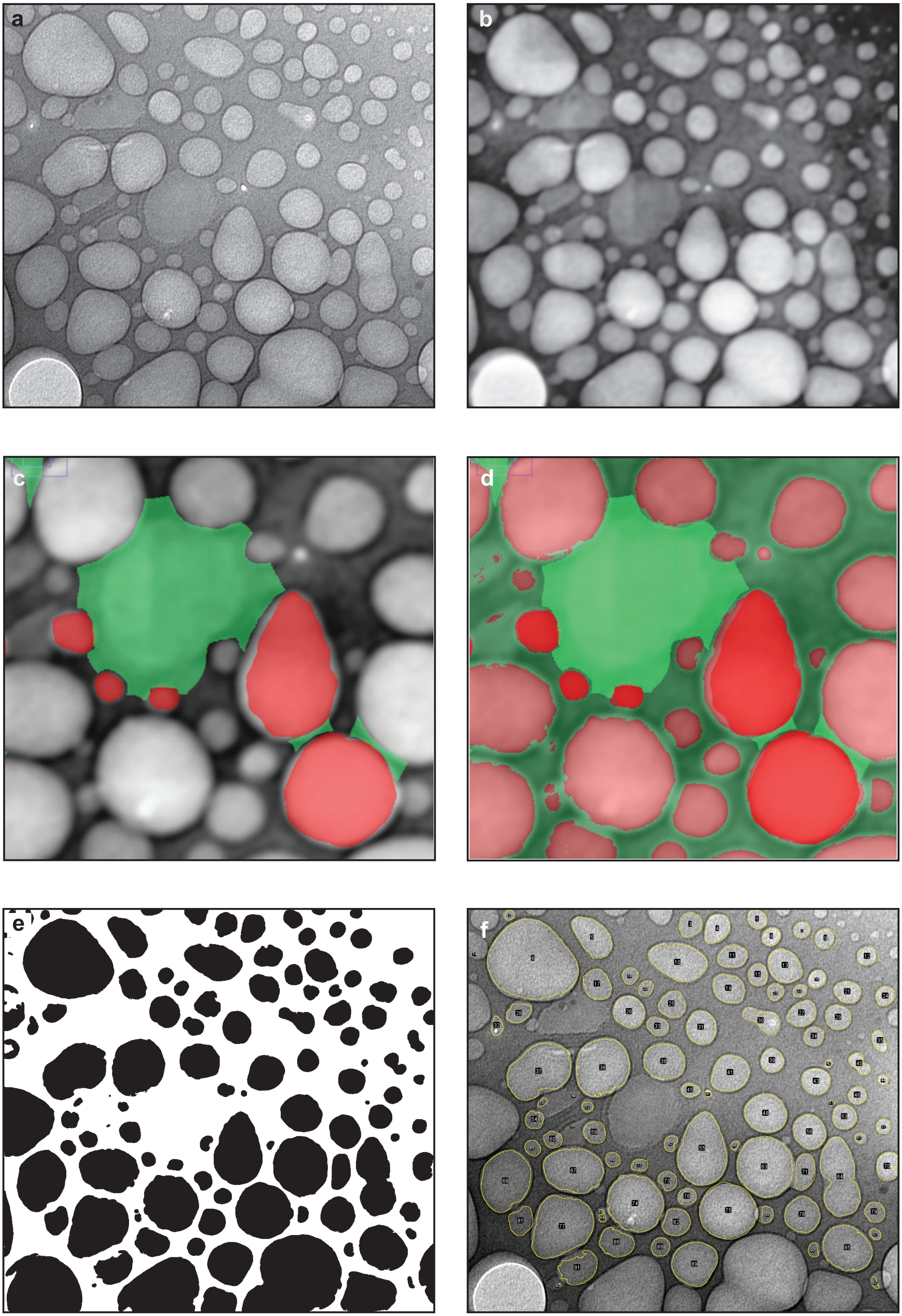

### S3

**Figure S3 Cyanobacteria Synechocystis sp. (PCC6803)**

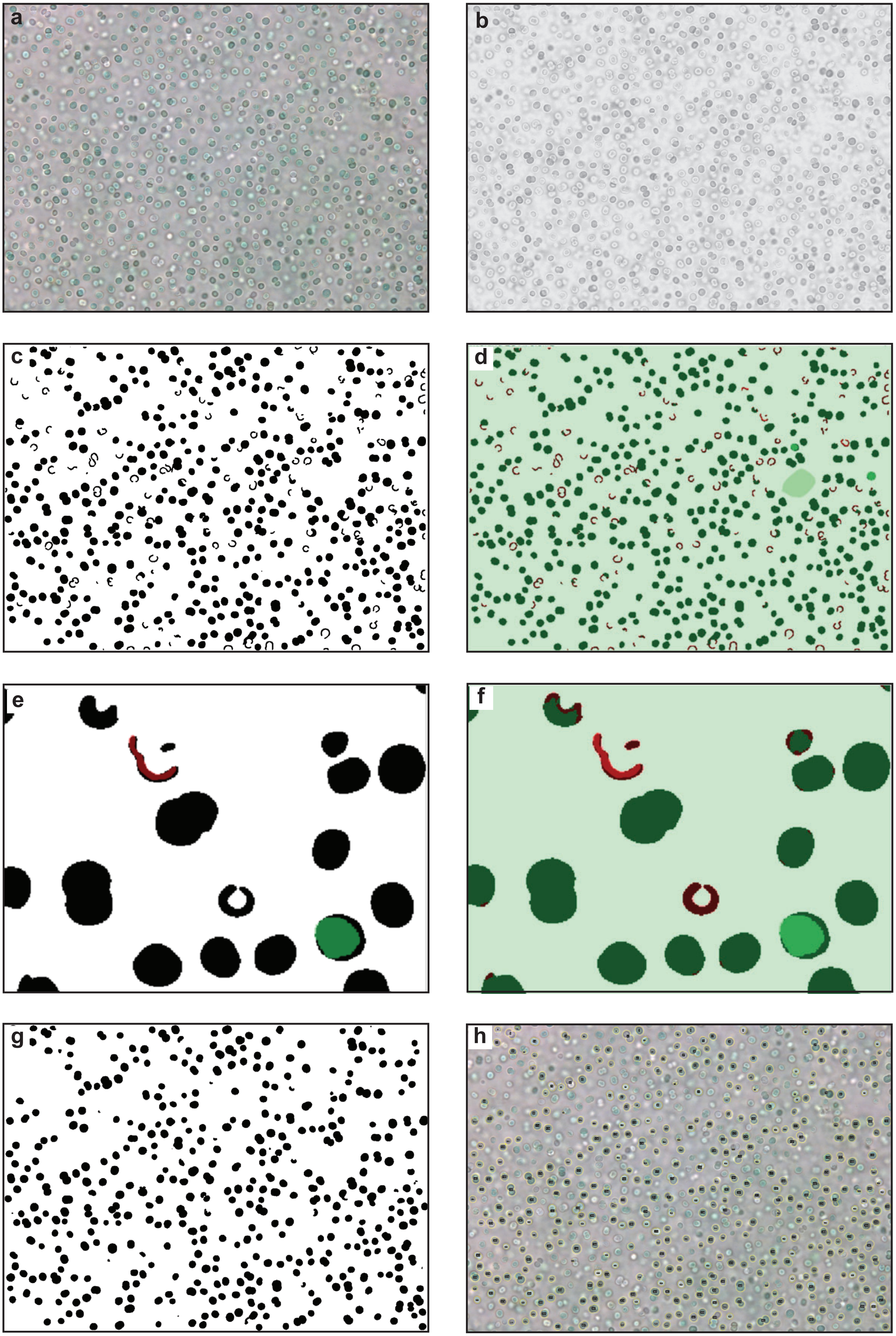

### S4

**Figure S4 Scatter plot and histogram**

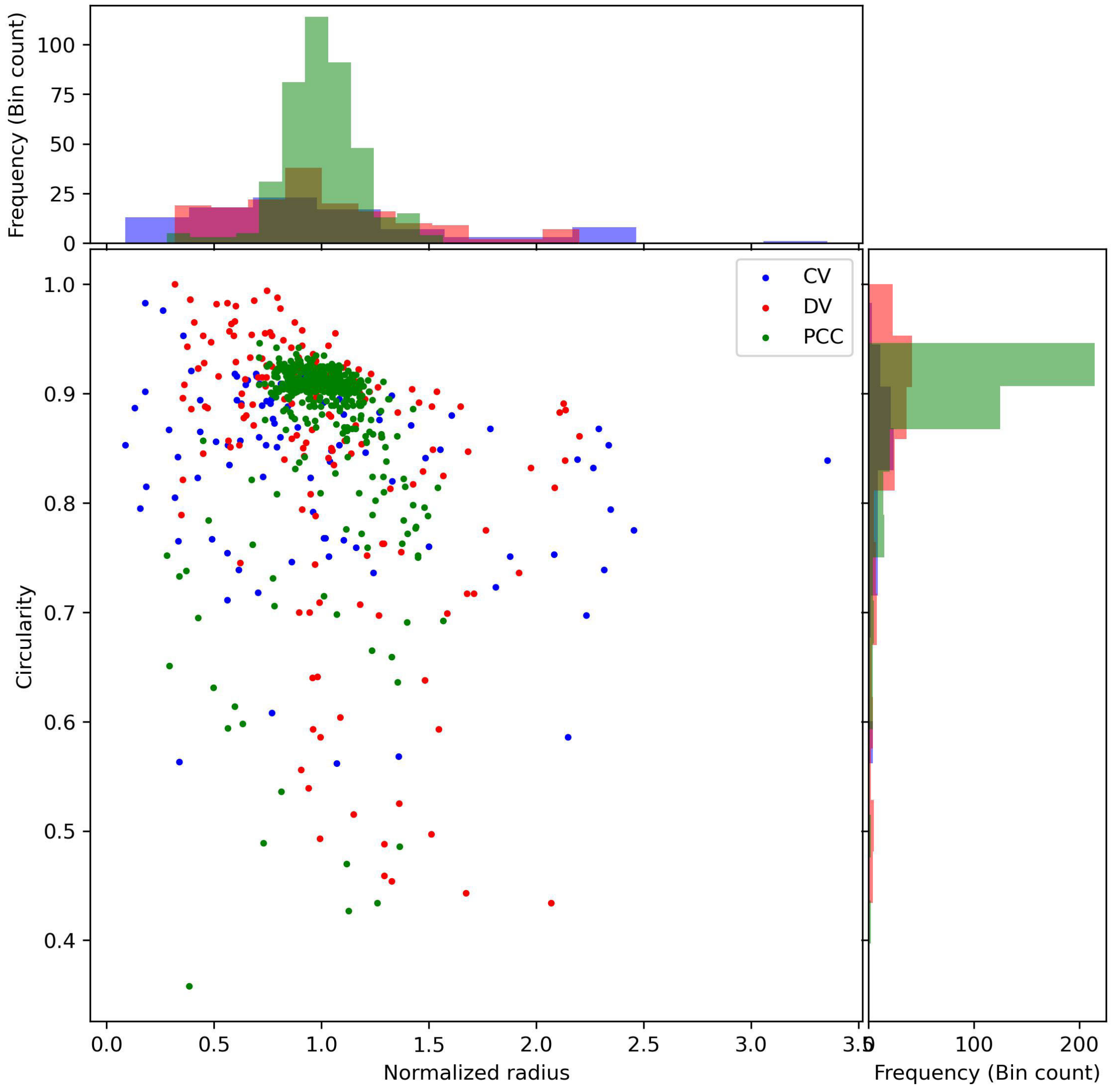
